# A Heuristic Derived from Analysis of the Ion Channel Structural Proteome Permits the Rapid Identification of Hydrophobic Gates

**DOI:** 10.1101/498386

**Authors:** Shanlin Rao, Gianni Klesse, Phillip J. Stansfeld, Stephen J. Tucker, Mark S.P. Sansom

**Affiliations:** Department of Biochemistry, University of Oxford, UK; Clarendon Laboratory, Department of Physics, University of Oxford, UK; OXION Initiative in Ion Channels and Disease, University of Oxford, UK

**Keywords:** Ion channel, hydrophobic gating, water, molecular dynamics, machine learning

## Abstract

Ion channel proteins control ionic flux across biological membranes through conformational changes in their transmembrane pores. An exponentially increasing number of channel structures captured in different conformational states are now being determined. However, these newly-resolved structures are commonly classified as either open or closed based solely on the physical dimensions of their pore and it is now known that more accurate annotation of their conductive state requires an additional assessment of the effect of pore hydrophobicity. A narrow hydrophobic gate region may disfavour liquid-phase water, leading to local de-wetting which will form an energetic barrier to water and ion permeation without steric occlusion of the pore. Here we quantify the combined influence of radius and hydrophobicity on pore de-wetting by applying molecular dynamics simulations and machine learning to nearly 200 ion channel structures. This allows us to propose a simple simulation-free heuristic model that rapidly and accurately predicts the presence of hydrophobic gates. This not only enables the functional annotation of new channel structures as soon as they are determined, but may also facilitate the design of novel nanopores controlled by hydrophobic gates.

**Significance statement:** Ion channels are nanoscale protein pores in cell membranes. An exponentially increasing number of structures for channels means that computational methods for predicting their functional state are needed. Hydrophobic gates in ion channels result in local de-wetting of pores which functionally closes them to water and ion permeation. We use simulations of water behaviour within nearly 200 different ion channel structures to explore how the radius and hydrophobicity of pores determine their hydration vs. de-wetting behaviour. Machine learning-assisted analysis of these simulations enables us to propose a simple model for this relationship. This allows us to present an easy method for the rapid prediction of the functional state of new channel structures as they emerge.

## Introduction

Ion channel proteins are water-filled, ion-conducting pores that are key components of biological membranes (1). In a resting or inactivated state, the flow of ions through the pore may be interrupted at one or more positions along the channel, termed gates. Upon activation, for example in response to ligand binding or a change in the membrane potential, conformational changes generally lead to expansion of the channel pore at its gate region(s) thereby switching it from a closed (i.e. non-conducting) to open (conductive) conformation. Ion channels represent attractive therapeutic targets and so there is considerable interest in elucidating the mechanisms underlying this process in many different ion channel families. In most cases, the determination of channel structures in various different conformational states provides the key molecular basis for understanding how this ‘gating’ is regulated, often supported by insights from electrophysiological and other biophysical approaches.

For a newly determined ion channel structure, its conductive state is most frequently inferred by measuring the physical dimensions of its transmembrane pore, displayed as a profile of pore radius along the central axis along with an image of the pore-lining surface. There are several methods for this, with one of the most widely used being the HOLE program (2). Such pore radius profiles provide an indication of the maximum size of ions that might be accommodated within a transmembrane pore, and steric constrictions narrower than the radius of a water molecule (∼0.15 nm) are considered as potential gates.

However, the permeation of ions and water through a given region in a sub-nanometre pore is influenced not only by its radius, but also by the local hydrophobicity of the pore lining. Ion permeation may readily occur through polar regions only just larger than the radius of the permeating species, but a hydrophobic pore segment of comparable dimensions may undergo spontaneous de-wetting to form a local nanoscale region devoid of water and ions (3–5). A particular channel conformation may therefore present an energetic barrier to permeation (i.e. be gated closed) in a hydrophobic region of the pore without requiring full steric occlusion. This is referred to as a hydrophobic gate (3, 6–8) or sometimes as a vapour lock (9). In such cases, the widening and consequent wetting (i.e. hydration) of the hydrophobic gate region enables the passage of ions through the channel (Fig. 1A).

**Figure 1:**
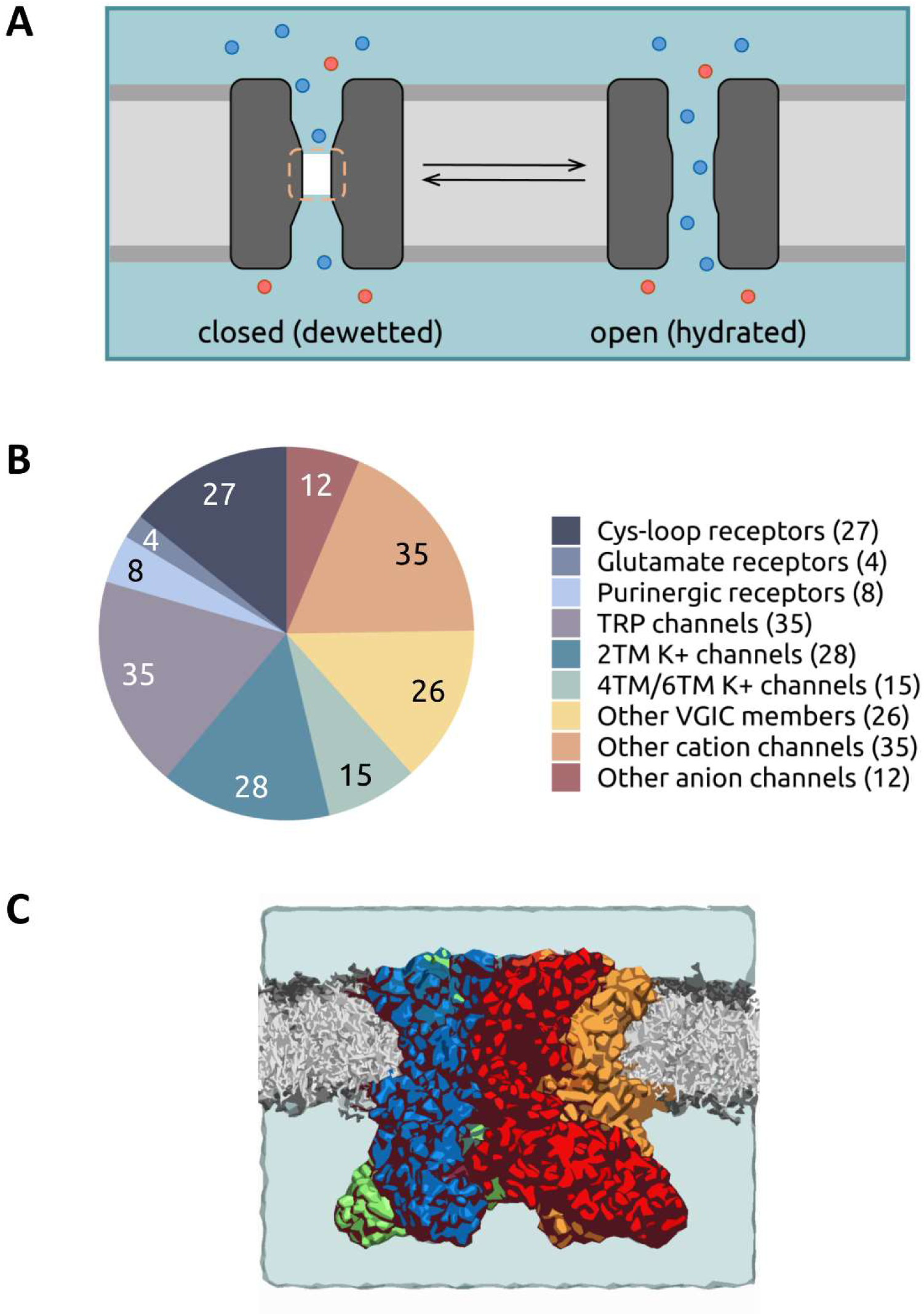
**A** Schematic of a hydrophobic gate in an ion channel. A hydrophobic gate region (dashed line) can spontaneously de-wet to form a dry (i.e. vapour) state and functionally close the channel. Widening of the gate leads to wetting (i.e. hydration) of the region and a functionally open state. Water is represented by the pale blue background shading, and ions by red and blue spheres. **B** Pie chart summarizing the ∼200 channel dataset which forms the basis of the current study. The channels are grouped into nine broad families: Cys-loop receptors, ionotropic glutamate receptors, purinergic receptors, transient receptor potential (TRP) channels, K^+^ channels (2TM, 4TM & 6TM), other members of the voltage-gated ion channel (VGIC) superfamily, other cation channels, and other anion channels. **C** Representation of a typical simulation system used in this study. Shown is the TRPV4 channel in a phospholipid (grey) bilayer. Water molecules and ions are present but omitted for clarity.

The concept of hydrophobic gating in members of the Cys-loop family of ligand-gated ion channels, including the nicotinic acetylcholine receptor (10), GLIC (7, 11), and the 5-HT_3_ receptor (12–14) is now well established. However, experimental and computational evidence of hydrophobic gates and barriers within other ion channels has also emerged, including for the TWIK-1 K2P channel (15), BK channels (16), the CorA magnesium channel (17), the CRAC channel Orai (18), and members of the transient receptor potential (TRP) channel family (19–22). Hydrophobic gating has also been demonstrated in synthetic nanopores (23, 24).

Computationally, molecular dynamics (MD) simulations of channel structures embedded within a lipid bilayer have aided the functional interpretation of new structures. Such simulations may range in complexity from characterisation of free energy landscapes of ion permeation and/or the conformational transitions associated with gating (6, 11) through to simpler simulations of water behaviour within channel pores (12).

The influence of pore radius and hydrophobicity on the wetting/de-wetting (i.e. the nanoscale liquid-vapour transition of water inside) of channels have previously been examined by simulation of model nanopores (3, 4, 25). For a uniformly hydrophobic constriction, the de-wetted (vapour) state appears stable below a critical pore radius of ∼0.5 nm, but this threshold radius decreases if the hydrophobicity of the pore lining is reduced (26). While channel structures containing hydrophobic regions with radii of ∼0.3 nm have been reported to represent de-wetted non-conductive conformations (e.g. (16)), our ability to predict this based on structure alone is hampered by a lack of detailed understanding of how pore de-wetting depends on the local radius and hydrophobicity, especially in complex biological structures such as ion channels where a wide range of hydrophobicity profiles are possible via different combinations of pore-lining side chains.

Fortunately, such analysis is now possible due to the recent explosion in structural data available for ion channels. This has mostly arisen from advances in structural biology, especially cryo-electron microscopy (27) and there are now >800 structures of over 100 unique ion channel proteins deposited in the PDB (Fig. 1B). Furthermore, improved software (e.g. CHAP; www.channotation.org) is now available for the analysis of channel pore dimensions, hydrophobicity profiles, and simulations of pore wetting/de-wetting (28). Thus, the ion channel structural proteome can now be subjected to a systematic examination of hydrophobic gating.

Here we use these improved approaches to quantify the influence of local pore radius and hydrophobicity on channel de-wetting via MD simulations of water behaviour in nearly 200 ion channel structures. The results of this systematic analysis provide a structure-based simulation-free heuristic model that allows rapid prediction of the conductive state of new channel structures as soon as they are determined. This method will also facilitate the design of novel nanopores (29) that contain hydrophobic gates (30, 31).

## Results

### A protocol for channel simulation and analysis

Each selected ion channel structure was embedded within a phospholipid bilayer, solvated on either side with 0.15 M NaCl (Fig. 1C), and subjected to three replicate 30 ns atomistic MD simulations to determine the behaviour of water within the transbilayer pore. Using our recently described channel annotation software (CHAP) (28), side chains that line the pore during simulations were identified and time-averaged profiles calculated for the pore radius, the local hydrophobicity, and the free energy of water as a function of position along the length of the pore (Fig. 2 and SI Fig. S1). Therefore, each point along the permeation pathway of a channel structure was associated with corresponding values for three variables of interest: pore radius, pore hydrophobicity, and free energy of water at that region of the pore. This enables the dependency of water free energy (and hence pore wetting/de-wetting) on local radius and hydrophobicity to be established and averaged across all the channel structures analysed.

**Figure 2:**
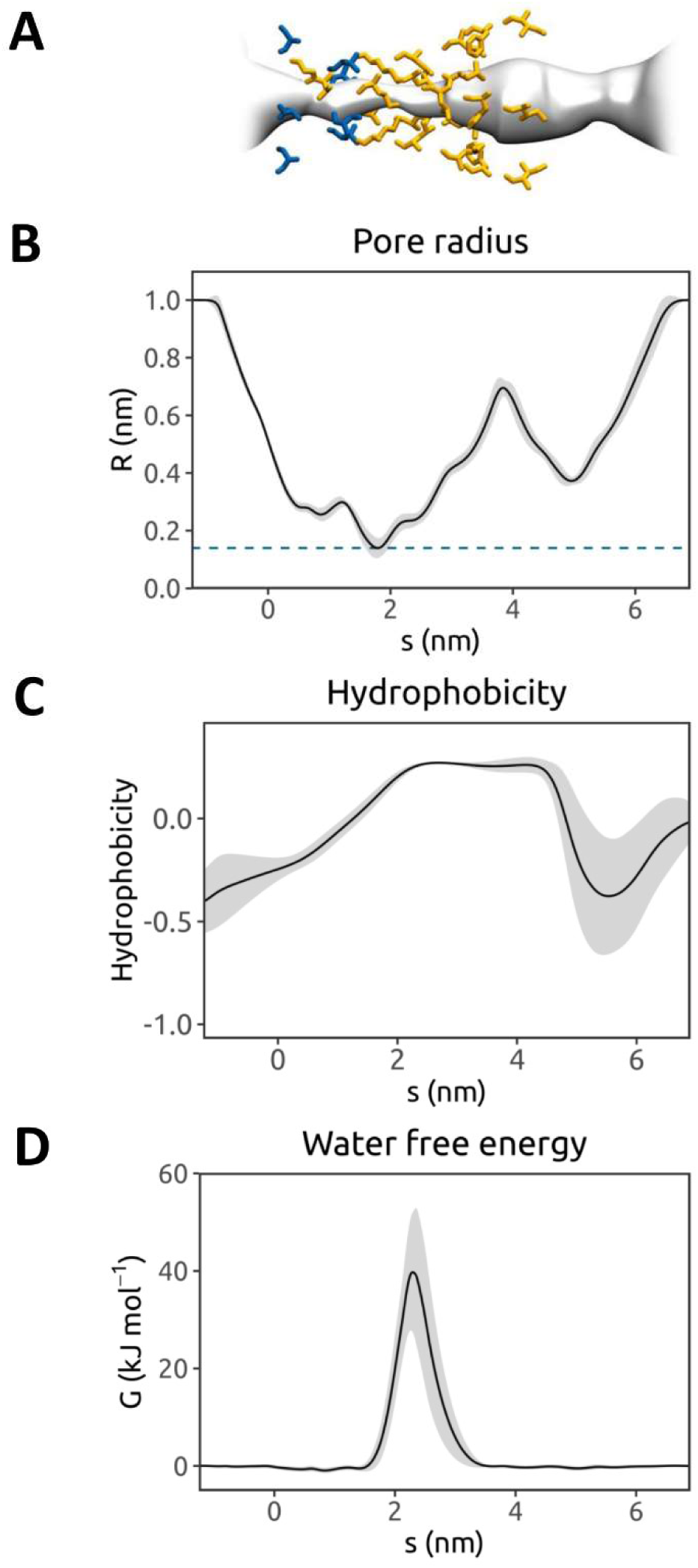
Annotation of an ion channel structure via MD simulations of water within the pore. This is illustrated for the TRPV4 ion channel (PDB ID 6BBJ; (46)). **A** Pore radius profile derived from one of 3 × 30 ns simulations of the protein embedded in a phospholipid bilayer. The mean radius, calculated using the final 20 ns of the trajectory with a sampling interval of 0.5 ns, is shown as a black line (with the grey band representing the standard deviation over time) as a function of position, s, along the pore axis. The pore-lining surface and the pore-lining residues of the channel (hydrophobic in orange; polar in blue) are shown at the top. **B** Hydrophobicity profile for the pore-lining side chains as a function of position along the pore axis s. The hydrophobicity values are evaluated using a normalised version of the Wimley-White scale (47). **C** Free energy for water as a function of position along the pore axis s. The free energy profile is obtained from the density of water within the pore as estimated from the MD simulation (see Methods for details).

During the simulations of water behaviour, positional restraints were applied to backbone atoms of the protein. This was to preserve the experimentally determined conformational state of the protein whilst allowing for local side chain flexibility. This is an important consideration as we are aiming to analyse water behaviour within known conformational states of ion channels, not to explore possible conformational changes of those channels by simulation, which would require much longer simulations. Nevertheless, we have previously shown that the presence/absence of such restraints does not significantly alter the free energy profile for water within a given simulation (32). Furthermore, we have also run more extended simulations to establish that 3 × 30 ns provides a robust estimate of the free energy landscape for water within an ion channel, even in the presence of a hydrophobic gate (SI Fig. S2). Indeed, tracking individual water molecules within a hydrophobic gate region which we hydrate at the start of the simulation reveals that de-wetting occurs within the first few nanoseconds (SI Fig. S3), as we have previously seen for model nanopores with hydrophobic gates (30, 33). Thus, for any individual channel our simulation protocol enables us to establish whether or not a hydrophobic gate is present, and the free energy profile for water within any such gate. Furthermore, the relative simplicity of the process allows us to perform simulation analysis across a dataset of nearly 200 channel structures that are representative of the ion channel structural proteome (SI Table S1).

### An ion channel dataset

To fully sample the range of radius-hydrophobicity combinations present within the ∼850 ion channel structures in the PDB, a reduced dataset was manually curated on the basis of structure quality (primarily resolution) and structural redundancy. Structures with a resolution of 5 Å or worse were excluded, as were those with an incomplete backbone trace in their pore-lining segments. Where several structures of the same ion channel species and of similar pore conformation (judged on root-mean-square-deviations and pore radius profiles along with visual inspection) were available, higher-resolution structures of the wild-type protein were chosen. This yielded a reduced dataset of ∼200 structures (see SI Table S1). The MD trajectory dataset (3 × 30 ns for each structure) formed the basis for subsequent analysis of pore de-wetting behaviour.

### Analysis of the main energetic barrier in each channel structure

Water free energy profiles derived from equilibrium simulations of wetting/de-wetting within a channel structure can be used as the basis for functional annotation (12). Any de-wetted region presenting an energetic barrier to water is likely to form a barrier to permeation (26), and thus water permeability may be used as a proxy for ion permeability. Based on our simulation dataset, channel structures containing one or more functionally closed gates were identified. These corresponded to ∼70% of the ∼200 structures analysed (SI Table S1). In Fig. 3A we show a point for each barrier to water permeability in the (*hydrophobicity, radius*) plane, coloured by the height of the water free energy barrier (*G*). Significant barriers (*G* > 2.5 kJ mol^−1^) were frequently associated with hydrophobic regions. The distribution of water free energy values on the pore hydrophobicity-radius landscape were also in qualitative agreement with predictions from simplified models of hydrophobic gating (4, 26), and in regions with elevated water free energies, hydrophobic aliphatic side chains (leucine, isoleucine, and valine) were found to be most prevalent (Fig. 3B).

**Figure 3:**
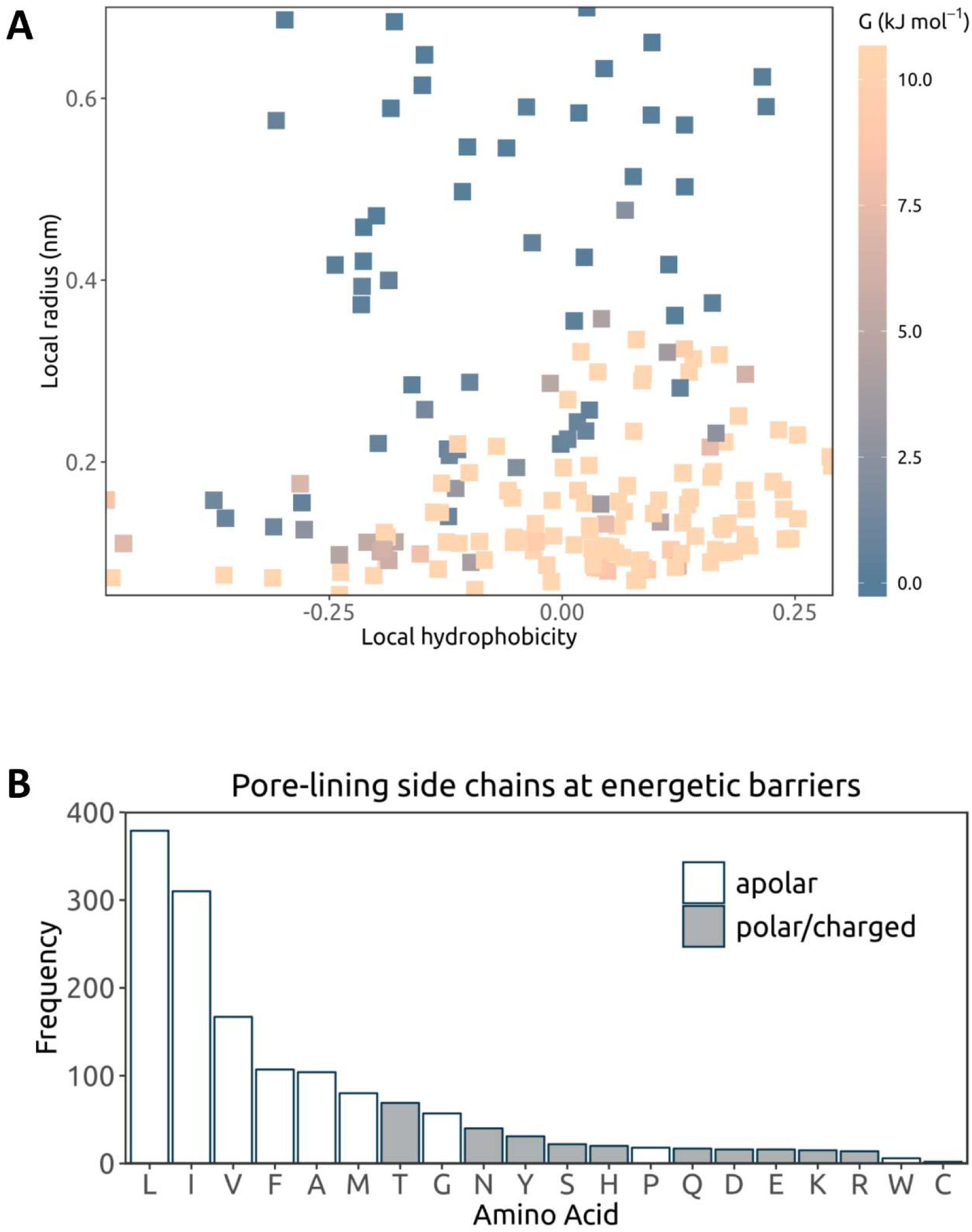
**A** Analysis of the main energetic barrier to water permeation in each ion channel structure. Each point corresponds to a single channel structure (see SI Table S1 for details), indicating the hydrophobicity and pore radius at the highest barrier in the water free energy profile (averaged across the three repeat simulations). The height of the barrier is given by the colour scale. Points with energy values outside the range are shown in the same colour as the lower or upper boundary. For structures containing a narrow pore-loop selectivity filter (e.g. K^+^ channels), the section on their water free energy profile corresponding to the filter region is excluded when locating the maximum energy point to represent the structure. **B** Frequency of the different pore-lining side chains at an energetic barrier in these ion channel structures, i.e. where the water free energy is greater than 1.5 RT (3.9 kJ mol^−1^) at the mean position of the residue along the pore axis s. Multiple amino acids may occur at each energetic barrier.

### Analysis of all residues lining ion channel pores

The relationship between local pore dimensions, hydrophobicity, and free energy of water at any given position along the channel axis also became clearer when all pore-lining residues in the ∼200 structures were surveyed, rather than just those present at barriers in the water free energy profiles. Examination of the distribution of water density values associated with all pore-lining residues revealed a substantive tail in the distribution corresponding to de-wetted regions, i.e. those with reduced water density relative to that of bulk water (SI Fig. S4). When a water free energy landscape in the local pore hydrophobicity-radius plane was constructed, a clear pattern emerged with the landscape clearly divided into two regions, corresponding to wetted and de-wetted pores (Fig. 4). In the hydrophobic region of the landscape an energetic barrier can be encountered in channel regions with radii up to ∼0.4 nm (the radius of a water molecule is ∼0.15 nm; Fig. 4). Conversely, hydrophilic regions of the landscape include pores which become hydrated at much smaller radii (i.e. < 0.2 nm). We note that the water free energy landscape is very similar even when alternative hydrophobicity scales are employed (SI Fig. S5).

**Figure 4:**
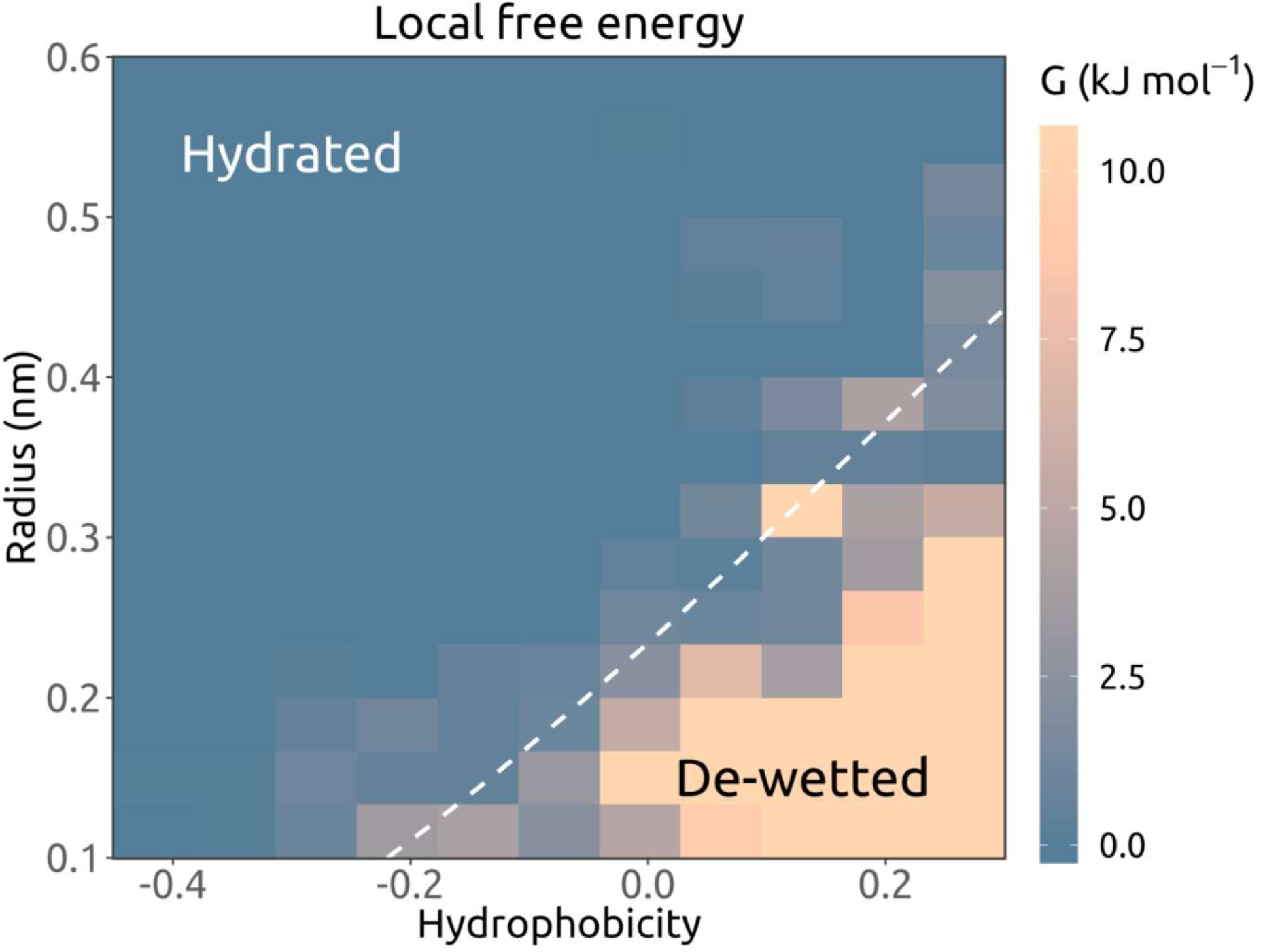
Analysis of all pore-lining residues for all simulated ion channel structures in terms of the local water free energy. From each simulation, mean measurements are noted at the mean position (*s*) along the pore axis (Fig. 2) of any side chain oriented towards the pore for at least 50% of the simulation. The local water free energy is shown as a function of hydrophobicity and pore radius for all occurrences of pore-lining side chains in the simulated structures. Hydrophobicity values are based on the Wimley-White scale, linearly normalised to [−1, 0.34], in relative units. The hydrophobicity-radius grid is coloured by mean energy. Regions with energy values outside the colour scale range are shown in the same colour as the lower or upper bound. The white dashed contour line indicates the 1 RT (2.6 kJ mol^−1^) position as given by a polynomial support vector machine classifier.

By training a support vector machine (SVM) classifier (34), we found that the hydrophobicity-radius landscape could be divided into two regions corresponding to the likelihood of pore wetting vs. de-wetting. These two regions are indicated by the dotted line in Fig. 4.

### A heuristic for predicting the conductive state

The water free energy landscape derived from the dataset of channel structures and simulations therefore allows us to devise a simple simulation-free heuristic technique for predicting the functional state of ion channel structures, based upon the correlation with pore dimensions and hydrophobicity alone (Fig. 5). Having used CHAP to identify pore-lining residues and to estimate pore radius and hydrophobicity profiles, the (*hydrophobicity, radius*) values for the pore-lining residues of a given channel are then mapped onto the landscape described above. By identifying the number of residues for which the corresponding (*hydrophobicity, radius*) points fall below the SVM classification line (i.e. in the de-wetted region of the landscape) the likelihood of a channel structure corresponding to a de-wetted and hence functionally closed state can be predicted. The sum of shortest distances (*Σd*) for residue points falling below the SVM open *vs*. closed classification line (corresponding to the dotted line in Fig. 4) provides a score indicating whether the channel structure is likely to contain a closed gate. As this machine learning-based predictive approach can be performed using automated analysis of a single set of coordinates (28) and does not require MD simulation of the structure, it can also be performed within a few seconds on e.g. a standard desktop.

**Figure 5:**
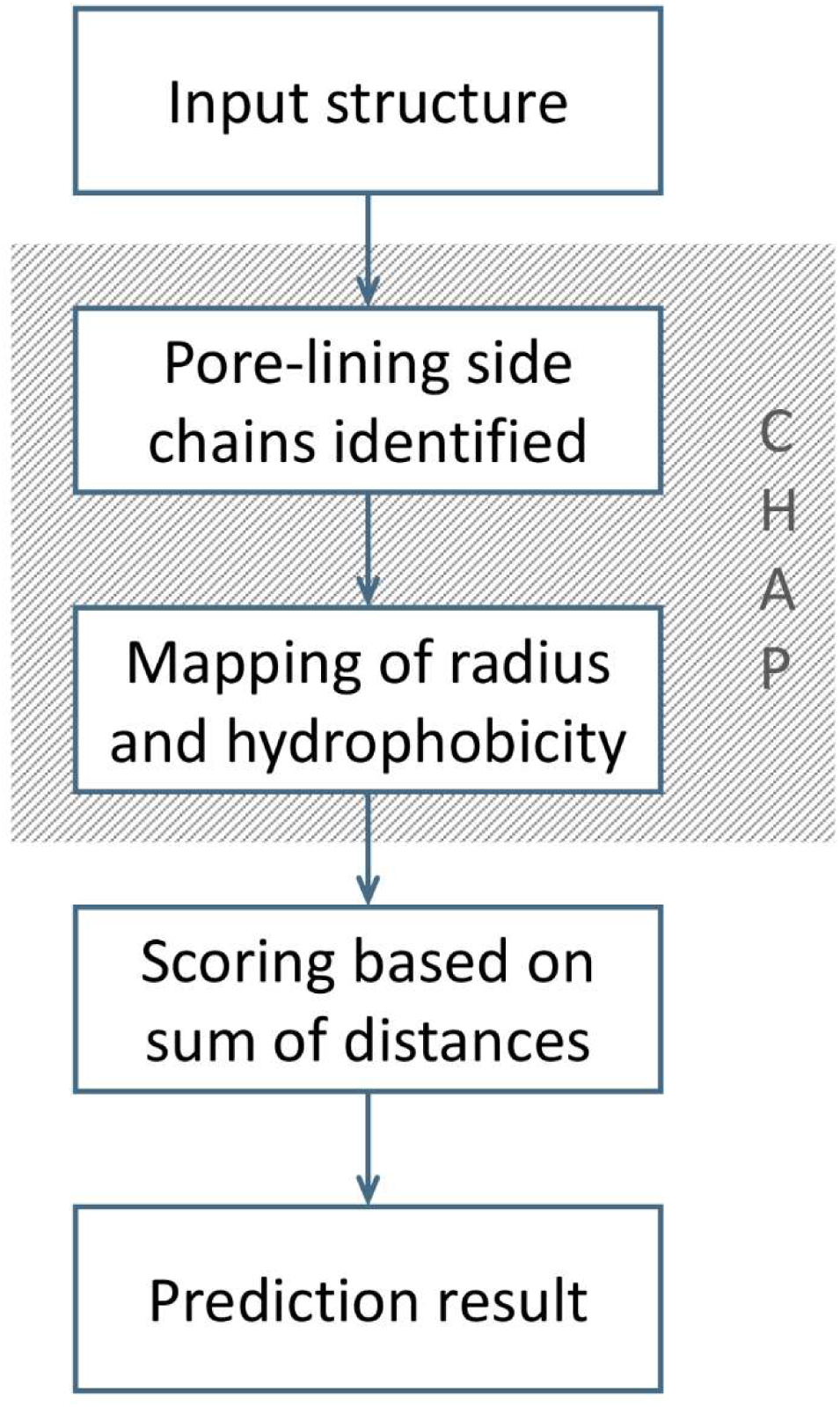
Schematic of a heuristic prediction approach for the permeation state of an ion channel structure based on analysis of hydrophobicity and pore radius profiles, both of which can be derived from a static structure using the CHAP analysis tool (28).

We illustrate the effectiveness of this heuristic approach using two recent structures (Fig. 6): one of the TRPV3 channel in a putative sensitised (but non-conductive) conformation (PDB ID: 6MHS) and one of the CRAC channel (Orai) in an open conformation (PDB ID: 6BBF) (35). For the open-state Orai channel, none of the (*hydrophobicity, radius*) points fall below the SVM classification line and so the structure is predicted to be fully wetted and correspond to a functionally open state of the channel. By marked contrast, for the TRPV3 structure, 12 points were below the SVM line, and the sum of shortest distances for residue points falling below this line was *Σd* = 1.4, leading to clear prediction of a closed hydrophobic gate for this particular channel conformation. Consistent with these predictions, when these two structures were subjected to the complete MD simulation and analysis protocols described above, the resulting water free energy profiles (SI Figure S6) confirmed the predictions made by this heuristic model.

**Figure 6:**
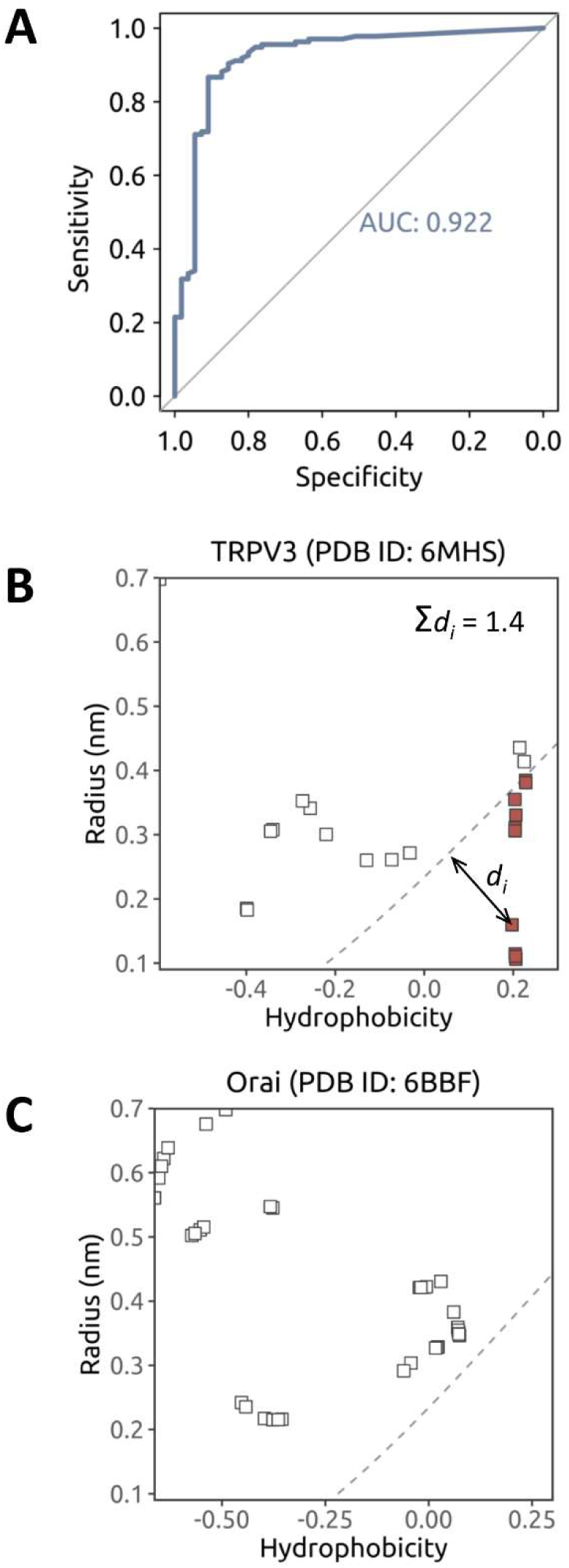
Illustration of the heuristic prediction approach. **A** ROC (receiver operating characteristics) curve analysis based on comparison of simulations of pore wettability (see SI Table S1) and heuristic based predictions indicating an optimal cut-off of *Σd* = 0.55 for the heuristic prediction of a closed channel, illustrated using two recent structures of (**B**) the TRPV3 channel in a non-conductive sensitised conformation (PDB ID: 6MHS) and (**C**) the CRAC channel Orai in an open conformation (PDB ID: 6BBF). Pore radius and hydrophobicity profiles are shown for the transmembrane domains of both structures. For each pore-lining side chain (represented as a coloured square on the pathway profiles), the channel pore radius at the residue is plotted against the corresponding hydrophobicity value. The sum of shortest distances between the dashed (1 RT) contour line and all points falling below it (coloured red), is then used as a score for identifying closed gates. A structure is predicted to be in a non-conductive state if it has a value of *Σd > 0.55*.

## Discussion

We have used MD simulations of pore water density to determine the wetting/de-wetting properties of nearly 200 ion channel structures that are representative of the currently available ion channel structural proteome. A systematic analysis of the behaviour of pore de-wetting within these structures as a function of local radius and hydrophobicity reveals an almost linear dependence of the critical radius for wetting upon the local hydrophobicity. Therefore, because pore radius and hydrophobicity profiles can now be readily estimated from any known structure, this analysis enables us to propose a simple simulation-free heuristic model for identifying closed gates in newly determined structures of ion channels or other forms of transmembrane pores (Fig. 7). This model is based on an underlying energy landscape derived from ∼600 simulations of water behaviour in ion channels, with a combined duration of ∼18 µs. It should be noted that pore wetting of a hydrophobic gate is necessary, but not always sufficient, for ion conduction to occur. However, our method provides a rapid and robust exploratory approach to the functional annotation of novel channel structures. Importantly, this analysis can be performed in a matter of seconds prior to more detailed simulations and/or experimental studies of the relationship between ion channel structure and function. Furthermore, this method may also facilitate design and engineering of novel nanopores (29) that contain switchable hydrophobic gates.

**Figure 7:**
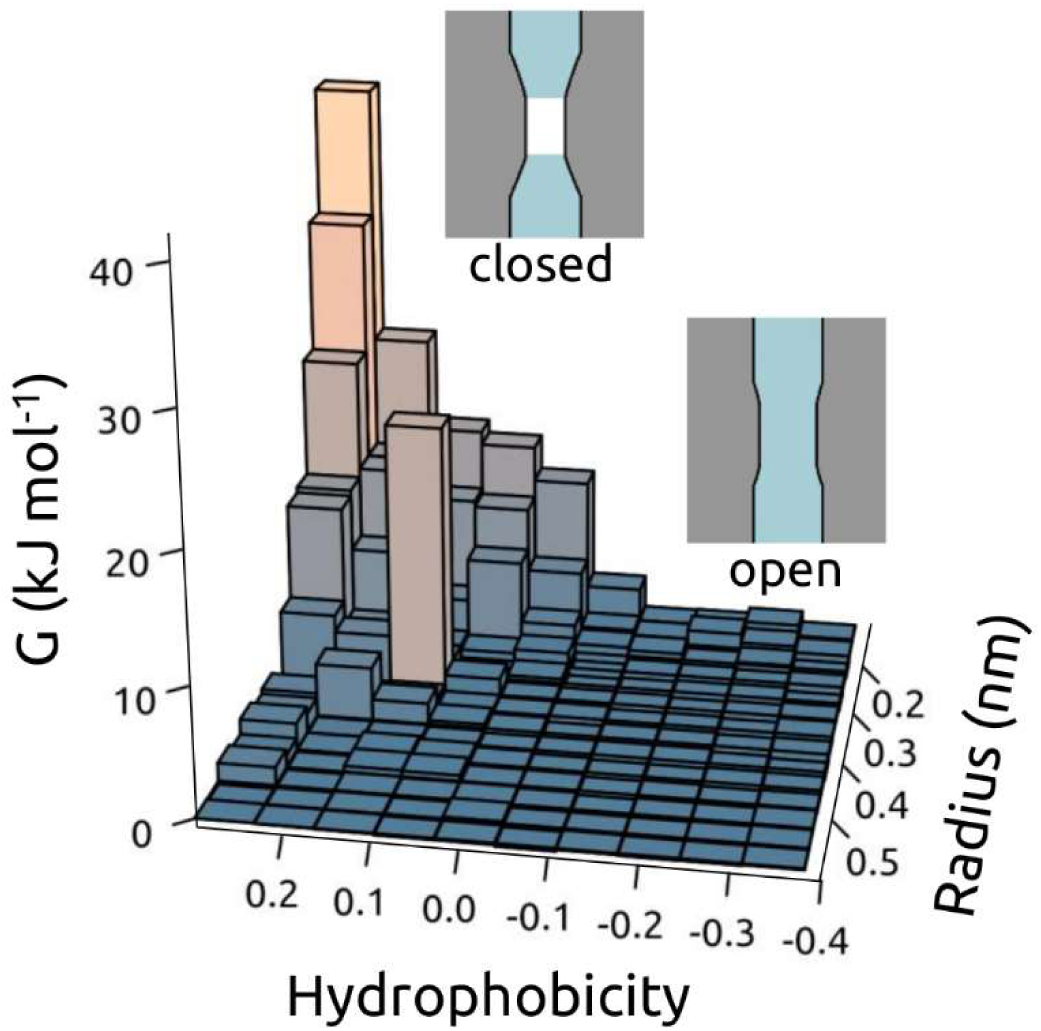
Schematic of hydrophobic gating as a function of (*hydrophobicity, radius*) of the transmembrane pore. The surface shows the free energy of water within a channel as a function of (*hydrophobicity, radius*) corresponding to the data in Fig. 5. Schematic depictions of de-wetted (closed) and hydrated (open) channel are shown for the two main regions of the data.

## Methods

The WHAT IF server (https://swift.cmbi.umcn.nl/whatif/) (36) was used to model incomplete side chains in the selected channel structures. Channel structures were inserted in a POPC (1-palmitoyl-2-oleoyl-*sn*-glycero-3-phosphocholine) bilayer using a multiscale protocol (37) for generating and equilibrating protein-membrane simulation systems. Molecular dynamics simulations were performed with GROMACS (http://www.gromacs.org/) version 5.1 (38), employing the OPLS all-atom protein force field with united-atom lipids (39) and the TIP4P water model (40). The integration time-step was 2 fs. A Verlet cut-off scheme was applied, and the Particle Mesh Ewald method (41) used to calculate long-range electrostatic interactions. Temperature and pressure were maintained at 37 °C and 1 bar, respectively, using the velocity-rescale thermostat (42) in combination with a semi-isotropic Parrinello and Rahman barostat (43), with coupling constants of τ_T_ = 0.1 ps and τ_P_ = 1 ps. Bonds were constrained using the LINCS algorithm (44). A harmonic restraint with a force constant of 1000 kJ mol^−1^ nm^−2^ was placed on the protein backbone atoms during simulations. The Channel Annotation Package (CHAP; www.channotation.org; (28)) was used to analyse trajectory frames, with bandwidths of 0.14 nm and 0.45 nm, respectively, applied for estimating water density and hydrophobicity along each channel axis. SVM classification was conducted using the Caret package (45) in R version 3.4.4 (www.r-project.org).

## Acknowledgements

This work was funded by BBSRC, EPSRC, the Leverhulme Trust, and Wellcome.

## Supplementary Information

**Table S1.**
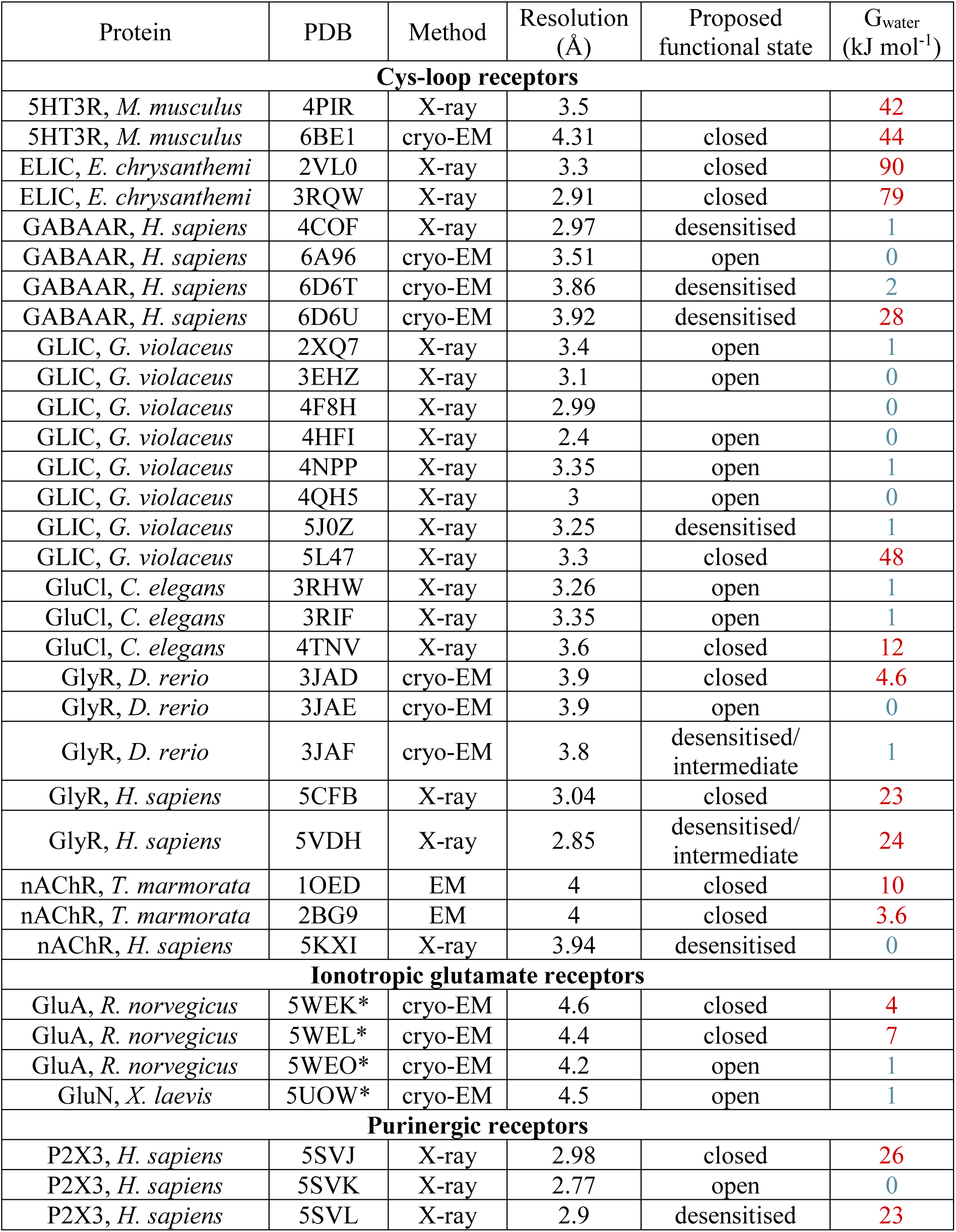
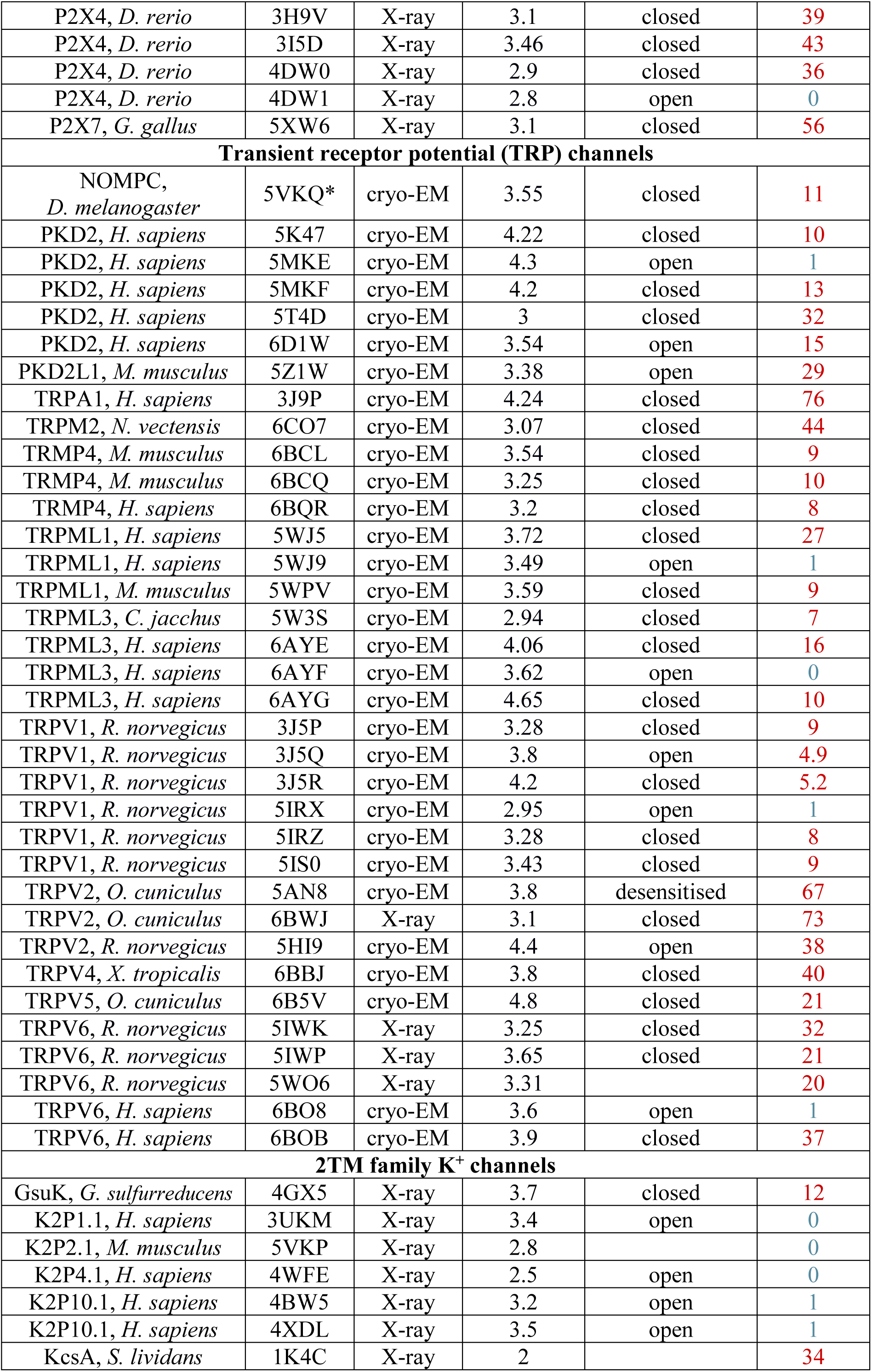
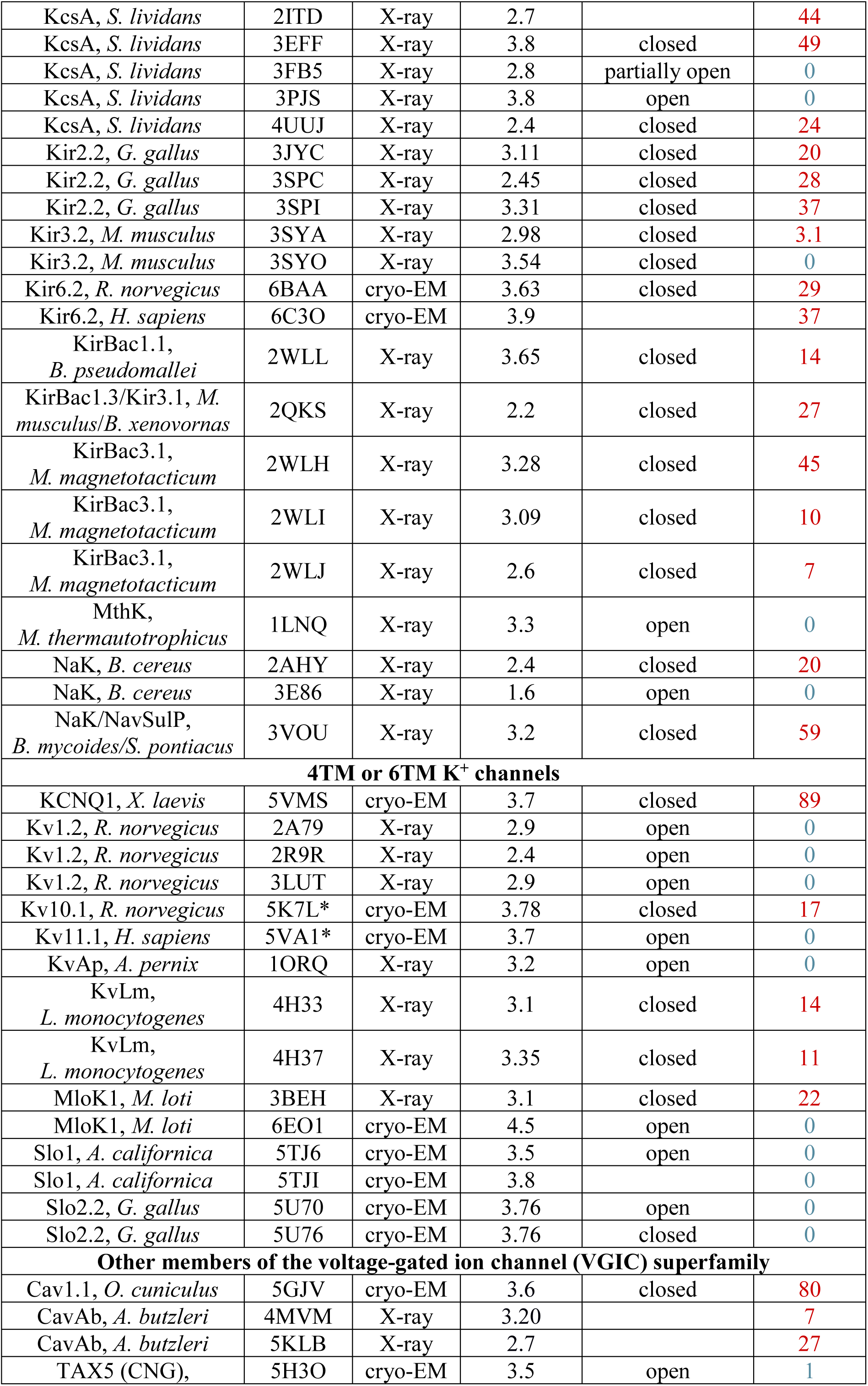
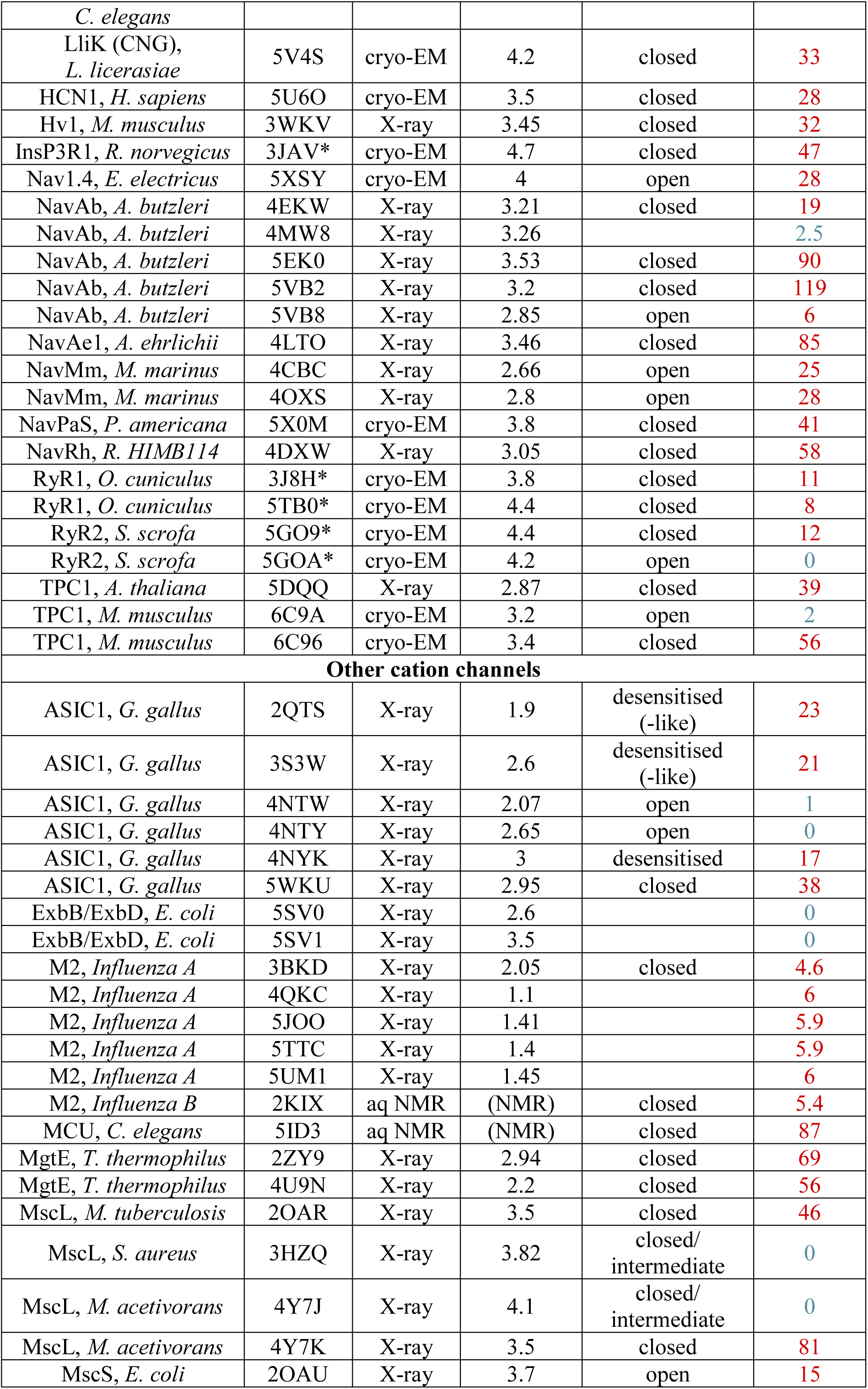
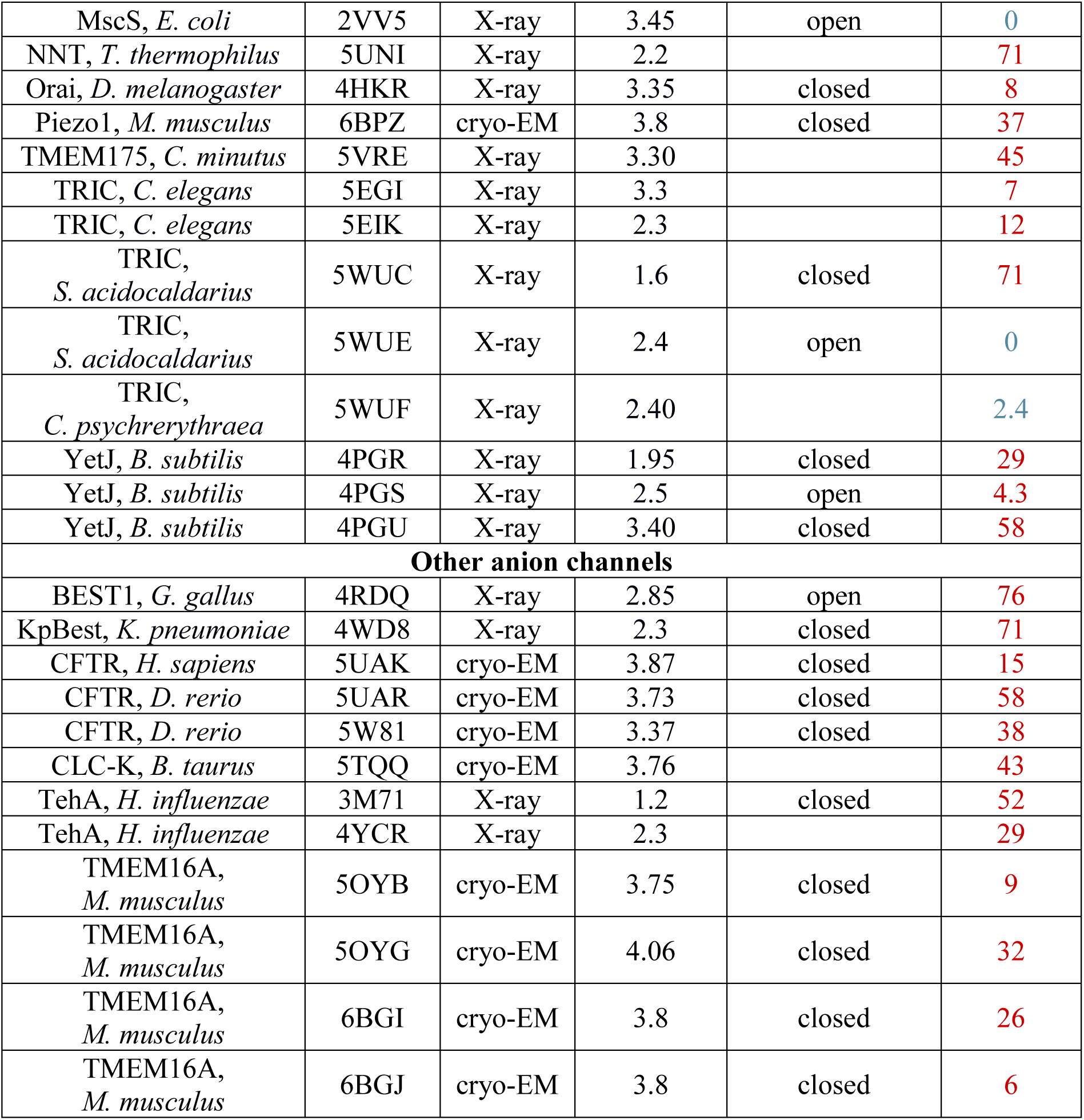
Ion channel structures analysed. Any auxiliary subunits, Fab fragments, ligands, ions, and/or water molecules present in a PDB entry were removed prior to molecular dynamics simulations. Large structures that were truncated to include only the transmembrane or pore domain for analysis are indicated by an asterisk (*) next to their PDB ID. Where available, the likely functional state of a channel structure as suggested upon its publication is noted. The height of the main energetic barrier to water (G_water_) is recorded for each structure, with values coloured red (> 1 RT) or blue (≤ 1 RT).

**Figure S1.**
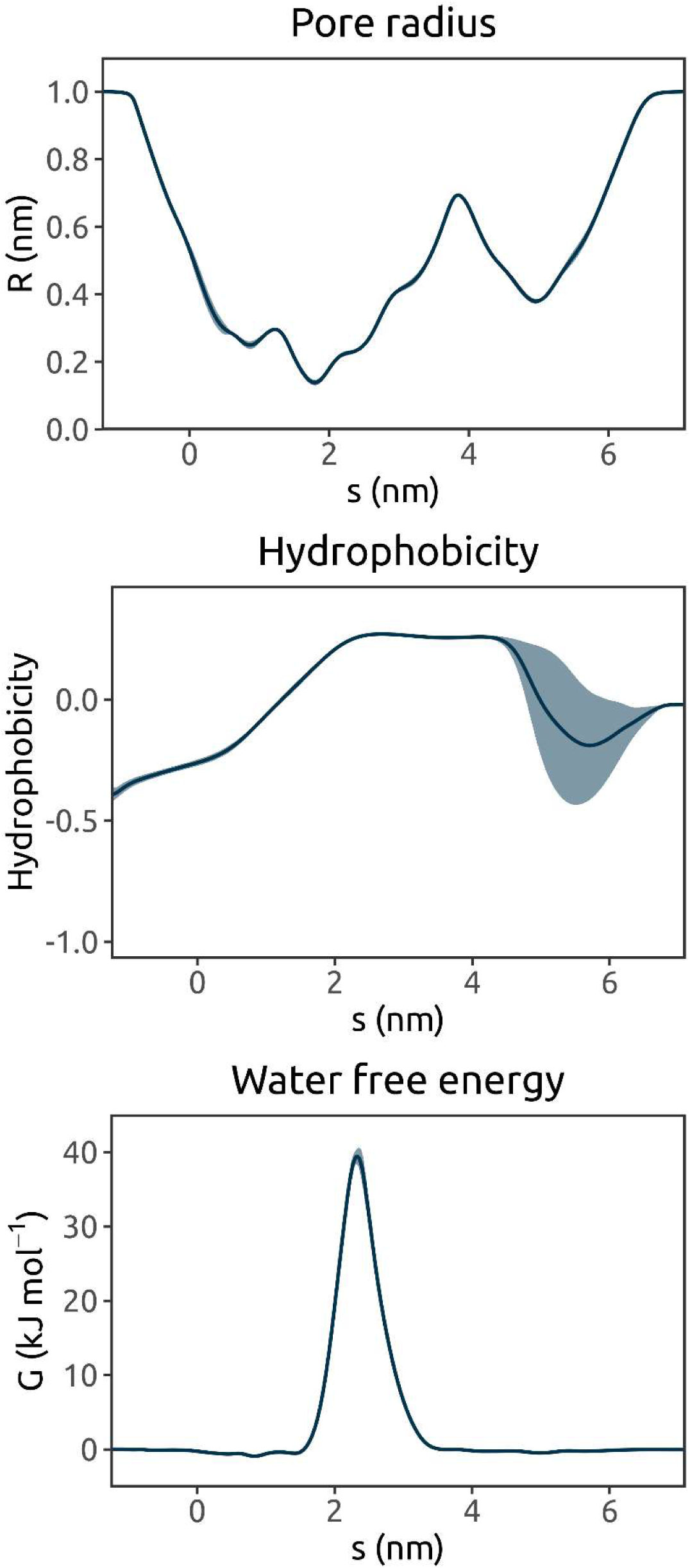
Average pore radius, hydrophobicity, and water free energy profiles derived from 3 × 30 ns MD simulations of water in the TRPV4 ion channel (PDB ID 6BBJ). The one-standard-deviation range is represented by the lighter-coloured band.

**Figure S2.**
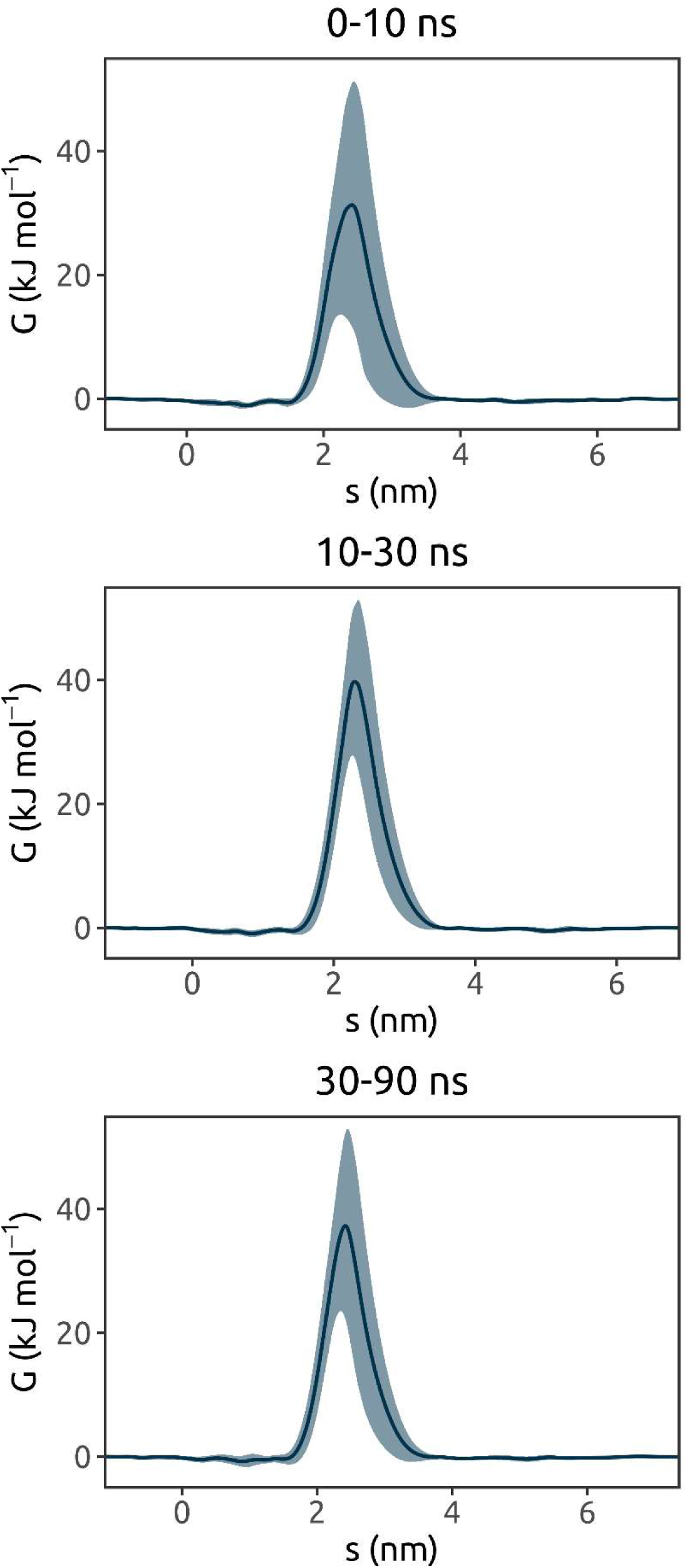
Water free energy profiles derived from different intervals of one 90 ns MD simulation of water in the TRPV4 ion channel (PDB ID 6BBJ), sampling every 0.5 ns. The one-standard-deviation range is represented by the lighter-coloured band.

**Figure S3.**
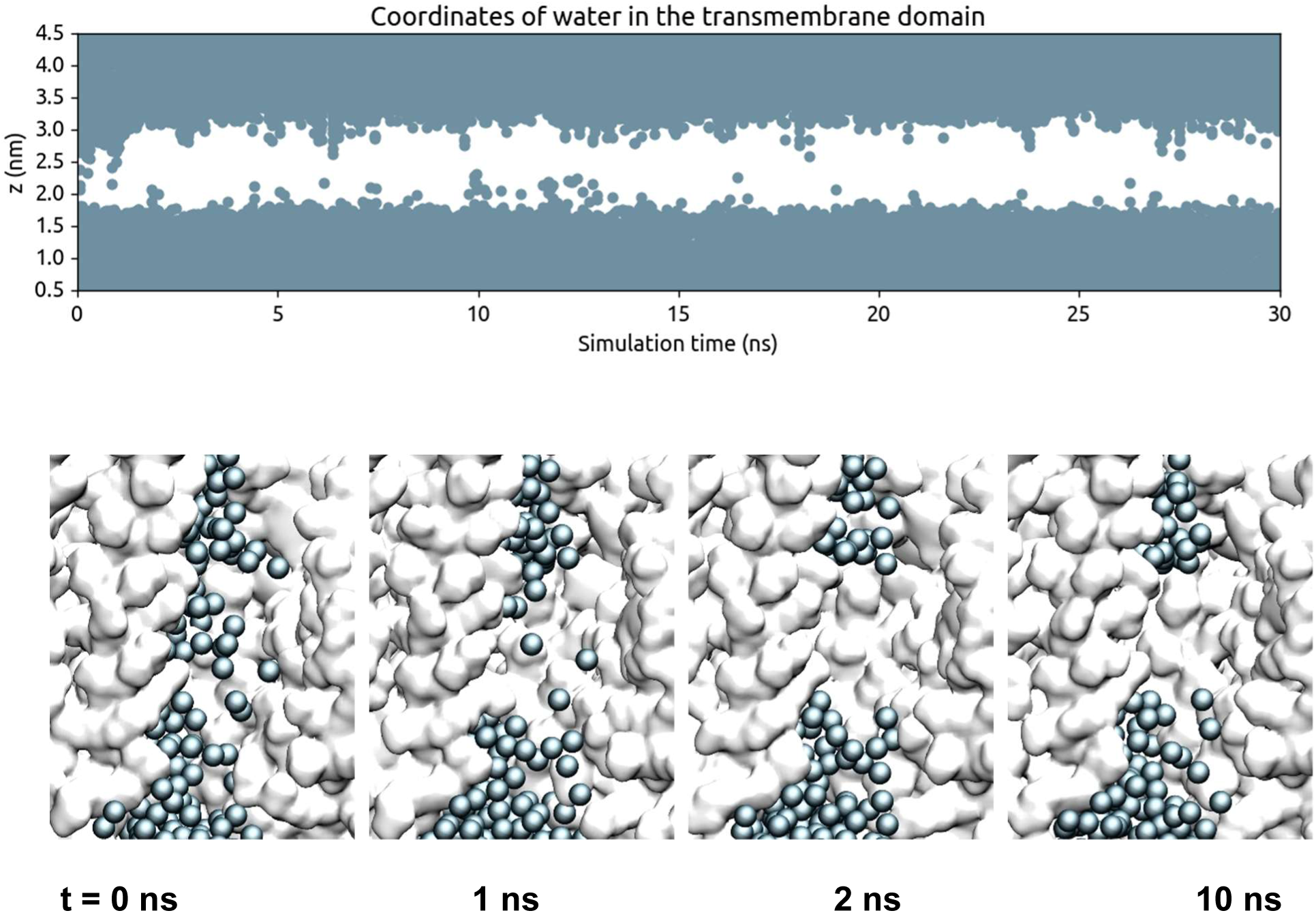
De-wetting of the hydrophobic gate region of the TRPV4 ion channel (PDB ID 6BBJ) in a 30 ns MD simulation. Water molecules are shown as blue spheres, with the protein in a surface representation.

**Figure S4.**
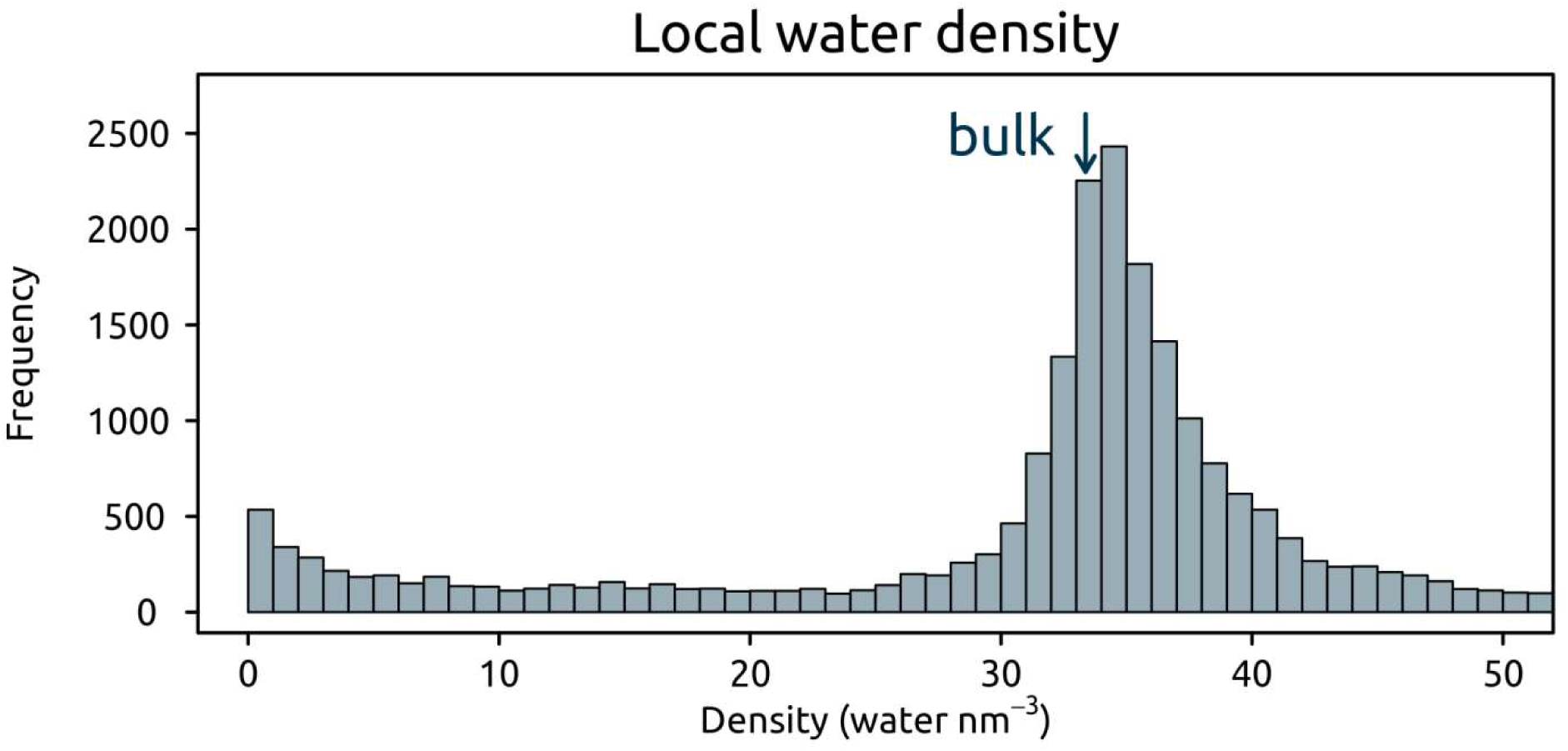
Distribution of local water density values for the dataset of all channel simulation, where the arrow indicates the density of bulk liquid water, at 33 water molecules nm^−3^.

**Figure S5.**
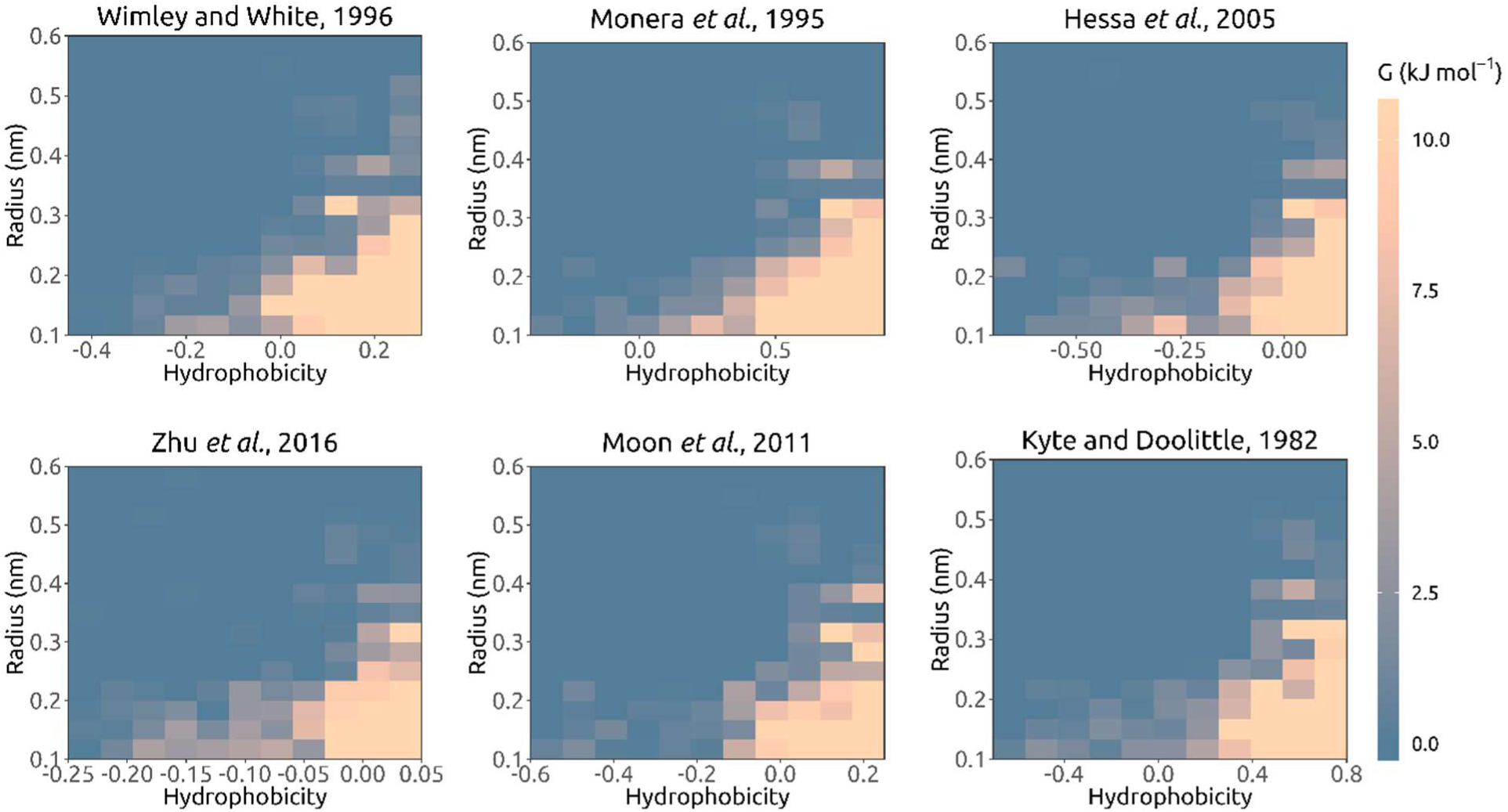
Local free energy as a function of hydrophobicity and pore radius, averaged over all occurrences of pore-lining side chains in the simulated channel structures. Alternative hydrophobicity scales are employed, each linearly normalized such that their respective positions of 0 hydrophobicity are unshifted and the largest absolute value amongst amino acids is equal to 1 (relative) unit, with more positive values indicating greater hydrophobicity. The scales compared are from: (1), (2), (3), (4), (5), and (6).

**Figure S6.**
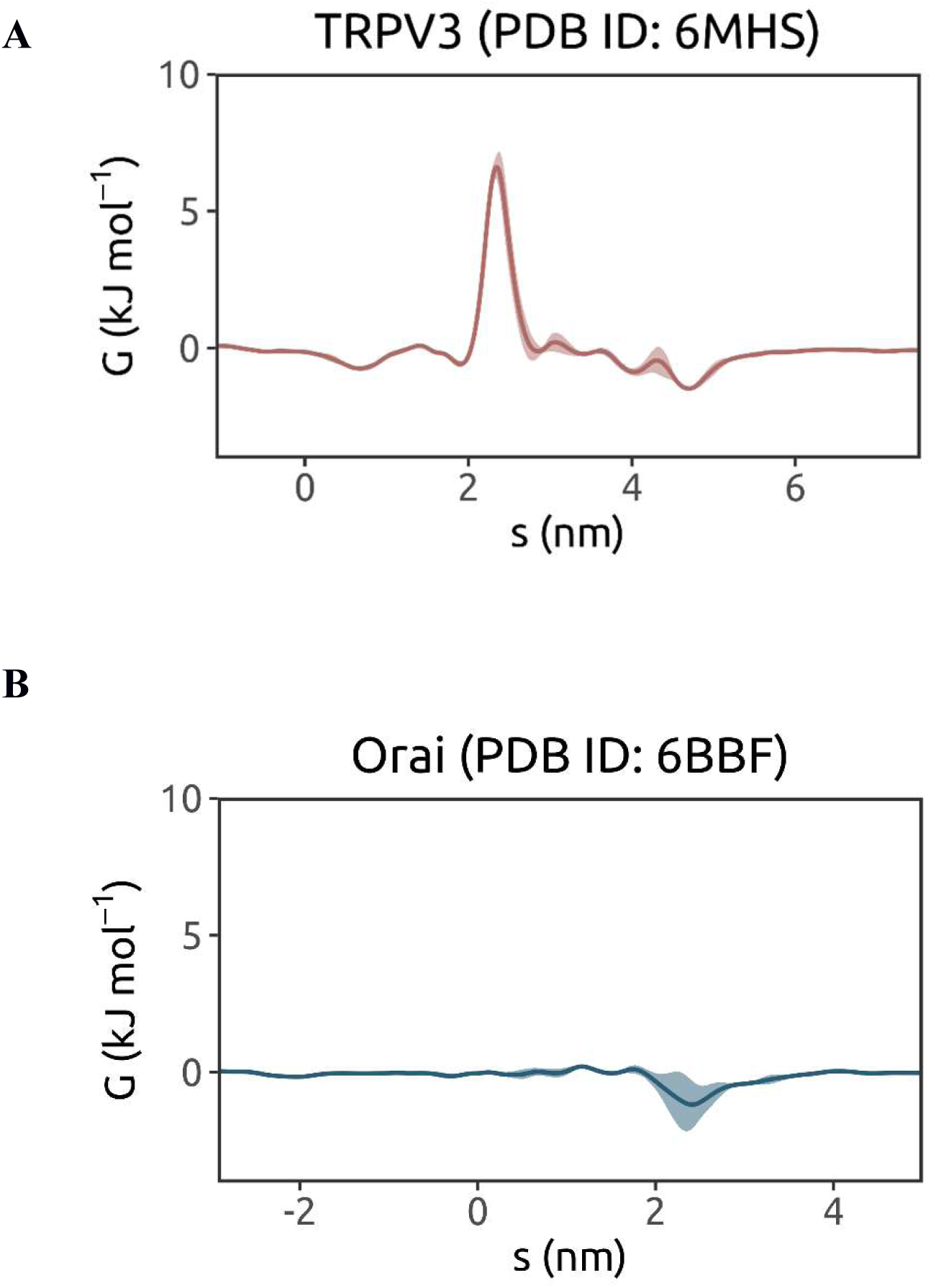
Water free energy profiles derived from triplicate 30 ns MD simulations of water of two recent structures of (**A**) the TRPV3 channel in a non-conductive (sensitized) conformation (PDB ID: 6MHS) and (**B**) the CRAC channel Orai in an open conformation (due to a H206A gain-of-function mutation; PDB ID: 6BBF).

